# Evidence for absence of bilateral transfer of olfactory learned information in *Apis dorsata* and *Apis mellifera*

**DOI:** 10.1101/473637

**Authors:** Meenakshi Vijaykumar, Sandhya Mogily, Aparna Dutta-Gupta, Joby Joseph

## Abstract

Capacity and condition under which lateral transfer of olfactory memory is possible in insects are still debated. Here we present evidence consistent with lack of ability to transfer olfactory associative memory in two species of honeybees, *Apis mellifera* and *Apis dorsata* in a PER associative conditioning paradigm where the untrained antenna is blocked by an insulating coat. We show that the olfactory system on each side of the bee can learn and retrieve independently and the retrieval using the antenna on the side contralateral to the trained one is not affected by the training. Recreating the paradigm in which the memory on the contralateral side has been reported at three hours after training we see that the memory is available on the contralateral side immediately after training and moreover, training with trained side antenna coated with insulator does not prevent learning, pointing to a possible insufficiency of block of odor stimuli in this paradigm. Bee does not learn the odor stimuli applied to one side alone as a stimulus different from odor presented to both sides. Moreover the behaviour of the bee as a whole can be predicted if the sides are assumed to learn and store independently and the organism as a whole is able to retrieve the memory if either of the sides have learned.

**Summary Statement:** The two halves of honeybee brain store and retrieve olfactory associative memories independently.

## Introduction

Lateral transfer of information helps environmental stimuli acquired, and learnt on one side to become accessible to both lobes of a bilateral brain (Aboitiz and Montiel, 2003; Gazzniga, 2000). This aids maximizing the computational ability of the brain by allowing each side of the brain to co-opt the other for joint decision or avoid duplicity of storage for efficient use of the substrate (Aboitiz and Montiel, 2003; Gazzaniga, 2000; Gazzaniga, 2014). Information transfer across the midline has also been theorised to be the basis of unified consciousness (Barron and Klein, 2016) and its importance has been highlighted in split-brain patients (Gazzaniga, 2014). In higher mammals, this crucial function is carried out by the *Corpus callosum*, a tissue present in eutherian mammals alone (Aboitiz and Montiel, 2003; Gazzaniga, 2000; Gazzaniga, 2014; Suarez et al.,2014). The formation of the corpus callosum has been suggested to be an evolutionary innovation (Mihrshahi, 2006), highlighting the importance of developing and evolving the process of transfer of information as an evolutionary stable strategy (ESS). Does this evolutionary jump have correlates in invertebrates such as insects? How important or necessary is it to have these bilateral connections in order to perform a task in invertebrates? Insects despite their primitive nature, are known to be able to perform complex tasks with their rather simple brains consisting of a few 100,000 neurons. Insect such as ants, wasps, honeybees, especially those belonging to the order *Hymenoptera* can perform complex tasks involving locating food sources, nesting sites and foraging back and forth between food source and nest, which would require coordination of vision, olfaction and navigation (Hansson and Stensmyr, 2011; Kaupp, 2010; Matsumoto et al., 2012; Roper et al., 2017; Sanes et al., 2010; Su et al., 2009). Bilateral transfer of information could aid the two lobes of the brain in coordinated decision making and also allow for one lobe to dominate in specialized tasks over the other, eg. speech, handedness in mammals, vision, prey escape and motor asymmetries in invertebrates (Anfora et al., 2011; Corballis, 2009; Corballis, 2017; Frasnelli, 2013; Frasnelli et al., 2014; Ghirlanda et al., 2009).

In free-flying bees (Masuhr et al., 1972) it was reported that side specific olfactory conditioning does not transfer to the contralateral side. It has been reported that in *Apis mellifera*, if the bee is trained in proboscis extension response (PER) to associate an odor with reward with stimuli applied to only one antenna, it can be retrieved by applying trained odor to the untrained contralateral antenna, three hours after training (Sandoz and Menzel, 2001; Strube-Bloss et al., 2016). In these experiments a wall was used to separate the two antennae and deliver the odor in a side-specific manner, arguing that the blocking the antenna using a coating influences the context of training and impairs transfer. In their study three hours post training, up to 50% of the bees responded by extending proboscis when the learned odor and not a novel odor was applied only to the contralateral antenna, suggesting the presence of a commissure relaying encoded odor specific memory between sides. In 2016, Guo. Y et.al reported the changes on a molecular level in the contralateral side after training even if the contralateral side was isolated by coating the antenna (Guo et al., 2016). This study, however, did not show transfer using behaviour, compared to controls trained with both sides closed. This group used silicon paste to block one antenna while training the exposed antenna to an odor. Post 24 hours the contra untrained antenna was checked for retention and transcriptomic analysis was carried out on the bees showing a change in the contralateral side over time. The results showed an up-regulation in memory and learning related genes on the untrained side of the brain, indicating a possible lateral transfer of this learned information and memory. Further, it has been recently reported that response of a subset of unidentified neurons called the Mushroom body output neurons (MBONs) which putatively receive input from the Kenyon cells (KCs) of the Mushroom body (MB) can change in a time dependent manner upon training to an olfactory stimulus (Strube-Bloss et al., 2016). Further, it was also reported that the memory transferred to the contralateral side to that of training is odor specific because they could show that the bees discriminated odors on the untrained side (Strube-Bloss et al., 2016). This would imply a bundle of fibres using population coding, or a small set of neurons using a complex temporal code connecting the two sides. Thus a plethora of evidence point to the possible presence of a commissure dedicated to the relaying of olfactory learned information from one brain lobe to the other. If this is true then recording the activity of the neurons in this commissure would also provide us insight into the nature of olfactory code, an exciting prospect.

The MBs are a crucial anatomical, higher processing centre of the insect nervous system that act as a multi-sensory integration unit (Strausfeld, 2002). The MBs have been suggested to play a central role in memory and retention (Menzel, 2012; Menzel and Benjamin 2012; Menzel and Muller, 1996). It has been posited that the MBs might play a role in the putative transfer of olfactory information from one lobe to another (Komischke et al., 2005; Malun et al., 2002). The area adjacent to the α-lobe has also been suggested to be the anatomical centres playing role in this transfer process (Komischke et al., 2005, Menzel and Benjamin, 2012; Menzel and Muller 1996; Okada et al., 2007, Sandoz and Menzel, 2001).

Work in our laboratory recently showed the presence of bilateral extrinsic neurons of the Mushroom body calyx (MB) in a species of grasshopper, *Hieroglyphus banian* (Singh and Joseph, 2018). In addition, a cluster of lateral horn (LH) neurons in *Schistocerca americana* have been shown to have a bilateral form bilateral innervation (Gupta and Stopfer, 2012). Very few correlates for lateral transfer of olfactory memory has been found other than the above-cited examples in insects. We therefore attempted to look for the neuronal basis of the phenomenon of bilateral transfer of information in a species of honey bee native to South East Asia, *Apis dorsata*, also referred to as the giant honey bee or the rock bee which is one of the crucial pollinators in the region, which is present only in the wild and hasn’t so far been domesticated. In our lab, olfactory pathway and PER conditioning in *Apis dorsata* has been shown to be very similar to *Apis mellifera* (Mogily et al., 2018). We trained *Apis dorsata* in PER conditioning to, pairing odor on one side with reward, while the contralateral side is closed with acrylic paint (Letzkus et al., 2006) and tested for retention on the contralateral side at 3 hrs post training and found no transfer. While testing the trained side antenna was closed with acrylic paint and the untrained side was open. Upon repeating this with *Apis mellifera* the results were consistent with our results in *Apis dorsata*. The learning rate and retention rate when both antennae are open is predictable by a model in which the bee decides to extend proboscis if either of the two sides decides to extend proboscis independently. To explain the discrepancy between these results and those from Sandoz and Menzel (2001) we repeated the procedure by Sandoz and Menzel (2001), using a partition of the same kind to prevent odor from reaching the untrained antenna and carried out two control experiments. One where we tested memory on the contralateral side immediately after training itself without a 3 hour delay and found it to be present. Second, even when the antennal on side being trained was covered with acrylic the bees learned when the isolation was attempted using the wall partition, indicating that wall is not an effective way for isolating one antenna from the other in our hands. These results from learning assays together with the absence of visible bilateral tracts between the olfactory pathways tract-tracing experiments (Mogily et al., 2018) force us to conclude that the olfactory pathways on the two sides of the brain learn independently and decides on the PER behaviour independently.

## Materials and methods

### Bee collection

*Apis dorsata* foragers were collected at 9 am from the flower sources such as *Turnura subtula, Tecoma stans, Eucalyptus globulus.* The bees were immobilized by cooling at 4ºC for ten minutes followed by mounting and tethering them in plastic holders using insulation tape. The bees were allowed to familiarize with this situation for two hours and then training was carried out. 15 minutes before training generic acrylic paint (Pebeo Studio Acrylics) was gently applied to one of the two antennae. Two control groups were always maintained during the training procedure, namely groups with both antenna open and both antennae blocked. Efficiency of the block was confirmed by the absence of learning in the group with antennae blocked and PER rates of this group was used as baseline for comparisons.

### Side-Specific Training for Apis dorsata

1-hexanol (Sigma Aldrich) was used to train the bees. Geraniol (Sigma Aldrich) was used to check for discrimination at 3 hours in *Apis mellifera*. Once the acrylic paint dried, the bees were divided into the three groups, one experimental and two control groups. Each bee placed on the pedestal for 14 seconds followed by the onset of the odor for 4 seconds (Conditioned stimulus-CS), the 30% sucrose reward (Unconditioned stimulus-US) was presented to the bee at the 3^rd^ second of odor onset and held for 3 seconds. (A 4 second CS and a 3 second US with 2 second overlap) Odor was delivered as a constant flow of air applied to the antenna via a 5mm diameter tube place 4cm away from the antennae. Odor was driven into the airstream from a 30ml glass bottle by pressurised air controlled by a valve. Glass bottles containing the aromatic liquid odors, were vacuum sealed and an odor delivery was carried out by a Teflon tubing connected to the glass bottle. A computer program controlled the valve and light emitting diodes that signalled the experimenter. In all the experiments, behaviour and physiology, an air suction exhaust was placed behind the animal so as to remove the odor after it had blown over the antennae. The bee would respond to the presence of the US by exhibiting PER (Bitterman et al., 1983; Matsumoto et al., 2012). A 10 minute inter-trial interval (ITI) was maintained between CS-US pairings and 5 trials were carried out with the entire training procedure lasting for one hour. The bees that spontaneously exhibited proboscis extensions were eliminated from the study. During the training if the bees extended their proboscis within 3 seconds of the odor onset (CS) they were counted as having odor evoked PER. For the bees that were trained with one antenna and checked with the same antenna (trained check), the acrylic coat was left intact on the untrained antenna. For the untrained test bees, the block was removed gently post training and the trained side was coated 15 minutes before testing. The schematic of the set of experiments is given in (Fig.1 A, B).

**Figure 1:**
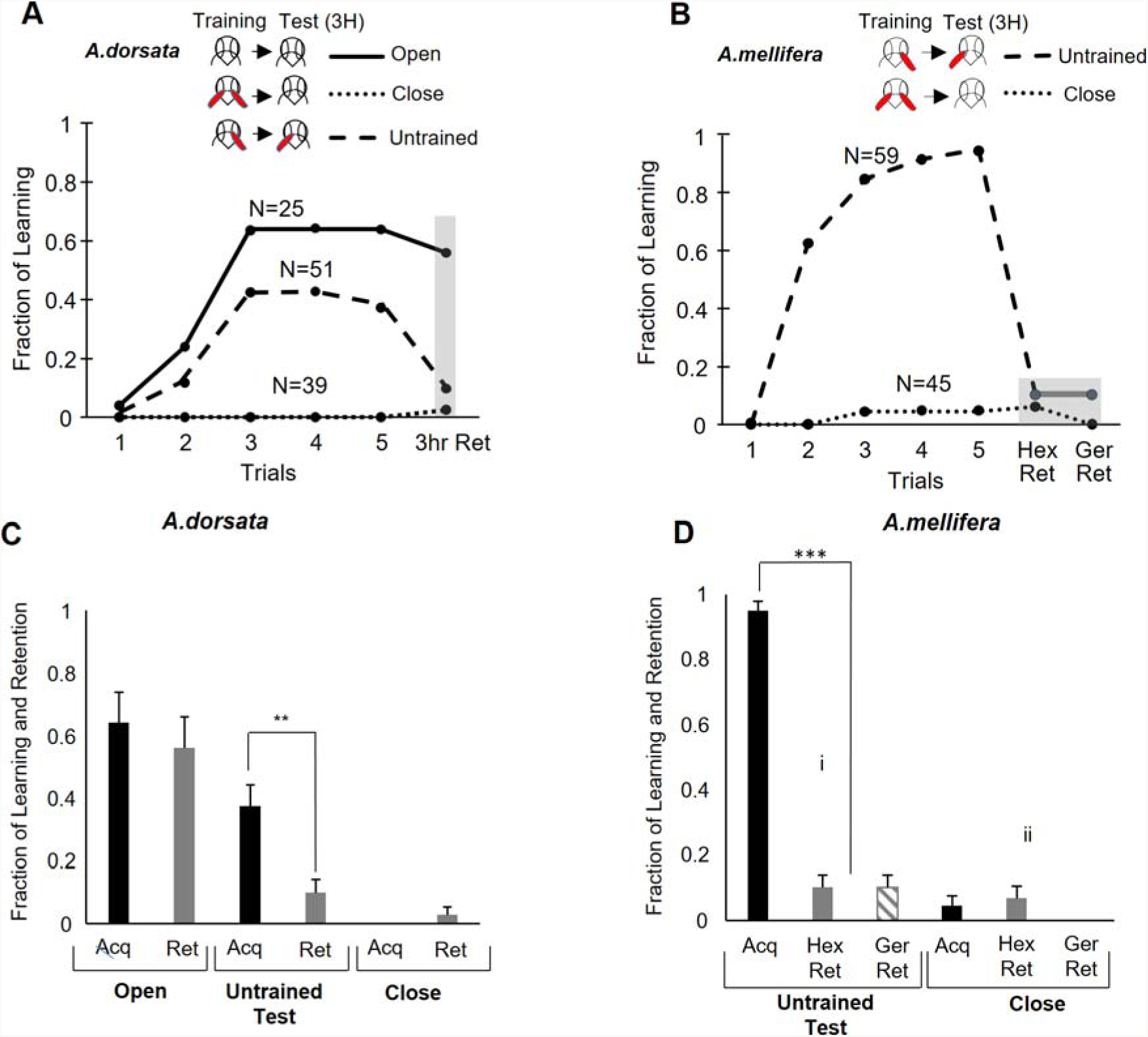
*Apis dorsata* and *Apis mellifera* do not show lateral transfer of olfactory memory. **A)** Testing for lateral transfer at 3 hours (3hr Ret) in *Apis dorsata* shows that memory on the untrained side at 3 hours is nearly zero (p=0.045 q=4). **B)** *Apis mellifera* bees also did not show any significant lateral transfer at 3 hours (p=0.22 q=1.5) though they had 95% acquisition. All the bees that responded to 1-hexanol (Hex Ret) responded to 1-geraniol (Ger Ret) on the transferred side at three hours indicating no discrimination (p=1). **C)** *Apis dorsata* showed significant difference between the learning (Acq) and retention (Ret) in the side contralateral to the trained, and no significant difference between the (i) untrained retention and (ii) closed antenna control group. The closed antenna group showed 0% learning and 2% retention which may indicate the success rate of our method of coating the antenna for blocking. **D)** *Apis mellifera* showed 95% acquisition but the transfer of memory to (i) contralateral side was similar to the group with (ii) both antenna closed.

To test that the acrylic paint was not causing damage to the antenna, in a group of *Apis dorsata*, the acrylic paint was applied to both the antennae and left for an hour (similar to the training period). The coat was then peeled off and the bees were trained and tested for PER conditioning (Bitterman et al., 1983). To confirm that the procedure of removing the paint was not causing a loss of memory by stress, a set of bees were first trained as per the one antenna blocked training protocol and 15 minutes before the retention test, a coat of acrylic paint was applied on both the antennae. Once dried, the coat was peeled off from the trained side antenna. The bees were then tested for retention to the odor memory. In all cases, identical protocols were followed for *Apis mellifera* in identical experiments.

### Checking for Contextual Stimulus

In order to confirm that coating with paint does not act as a contextual stimulus, *Apis mellifera* were first trained with either antenna covered with a coat. The trained bees were then divided equally into two groups, one set of bees were tested for retention at 3hrs with the coat on and the other set were tested for retention with the coat removed.

### Side-specific training with wall partition

We carried out the side-specific training for *Apis mellifera* using a wall barrier as specified in protocol in (Sandoz and Menzel, 2001). A plastic wall (40mm x 50mm) cut in the shape side profile of the bee; in its holder was used to separate the two antennae. The wall was placed such that the mandible and proboscis were adjusted slightly to one side depending on which antenna was to be trained-lobe. The spaces between the wall and bee’s head were sealed with wax. An exhaust vent behind the setup constantly drew air peeling away from the preparation (Supplementary Fig.1).

## Results

### No lateral transfer of memory in *Apis dorsata*, (A+/0)

The learning rate for bees with one antenna blocked at the end of the 5^th^ trial reached 37% (n=52) (Fig.1 A, C). The learning and retention with both antennae closed was negligible as expected. The retention test with the untrained antenna was not significant compared to the condition where both the antenna were closed (p=0.045 Cochran’s q=4) consistent with the absence of lateral transfer of memory. We corrected the value of α using Bonferroni correction to a value of α=0.044.

### No transfer or discrimination in *Apis mellifera*, (A+/0)

For Apis *mellifera* bees the acquisition reached 95% (n = 59) at the end of the 5^th^ trial (Fig.1 B, D). Given the high learning and acquisition rate, only the, both antennae closed control group was maintained through the training procedures. The learning rate in the contra test group was nearly zero and not different from both antenna closed group (N=35, p=0.22 q=1.5). The test for retention on the untrained side showed a significant drop in the percentage of retention (10.5%), this value was close to the retention of the closed antenna control group (6.8%). There was also no odor discrimination exhibited by the bees which had contra retention, the bees which responded to 1-Hexanol also responded to 1-Geraniol (Fig.1 B, D) while those with both antennae open (n=52) showed good discrimination between Hexanol and Geraniol.

### Performance in learning and retention were consistent with the olfactory pathways in the two sides acting independently

If decision by any of the two sides can cause PER, then one would expect that the probability of evoking PER should be predictable from the learning rate and retention rates of one side alone. Sum of probabilities of either of them deciding to evoke PER minus the probability that both of them would. For the learning rate, the prediction would be (2*0.38-0.38*0.38=0.62), approximately equals 0.64, the observed learning rate. Same should follow for retention (2*0.35-0.35*0.35=0.57) approximately equal 0.56, the observed retention rate. These predictions match, indicating that the two sides make decisions independently and indicates no lateral transfer of olfactory learnt memory, while learning or after 3 hours.

### *Apis dorsata* showed high memory retention on trained side, be it left or right, (A+/0)

The learning rate for bees with one antenna blocked at the end of the 5^th^ trial reached 38% (n=34) (Figure 2 A, B). The learning rate of the bees with both antenna open reached 64% (n=25) at the end of the 5^th^ trial and bees with both antennae closed showed 0% for n=39 learning. 92% of the bees that learned retentained on the trained side at 3 hours. Open antenna bees exhibited 87% retention and a 2% retention was seen in bees with both the antennae closed. The difference in retention rate between the one antenna trained and tested group and both antennae trained and tested control group was found to be insignificant (p=0.13, Cochran’s q= 5.77). A significant difference was found between the retention of the trained tested and the group of closed antennae bees (p=8×10^−4^, Cochran’s q= 11.15) (Figure 2 A). No significant difference was seen between the acquisition and retention rates of left and right antenna trained bees.

**Figure 2:**
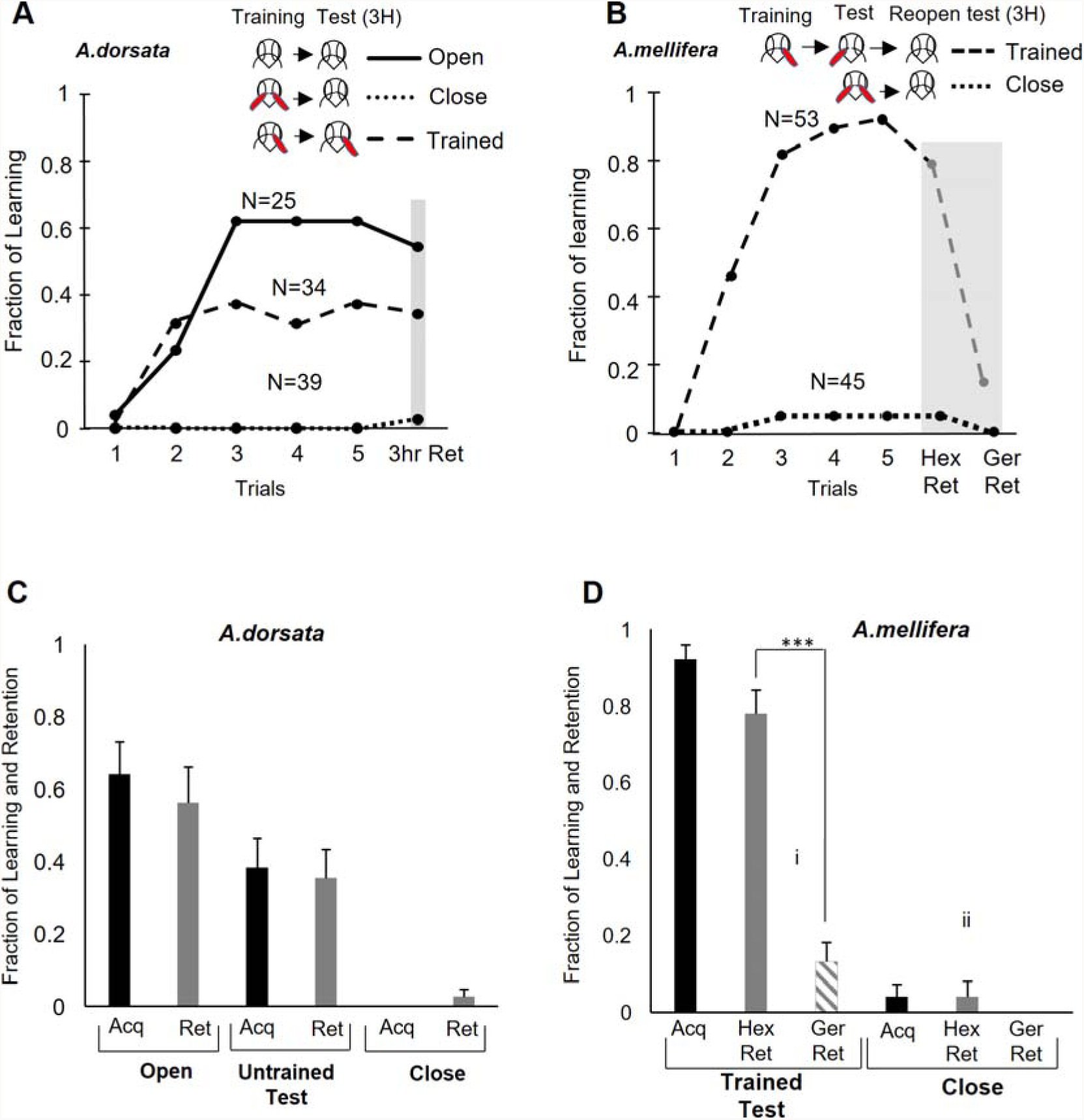
Memory is retained on the trained side in *Apis dorsata* and *Apis mellifera*. **A)** *A. dorsata* learned and retained memory after 3 hours (3hr Ret) with one antenna if tested with the same antenna and also learned and retained if trained and tested with two antennae. The bees showed significantly lower acquisition and retention with one antenna compared to two antenna training. **B)** For *Apis mellifera* coating and removing the coat does not affect the acquired memory on the trained side, difference between the retention of Trained tested and closed control group was significant (p=8×10^−4^, q=11.15). *Apis mellifera* showed 95% acquisition, and retention to 1-Hexanol was seen to be stable once the trained ipsi antenna was unblocked at 3 hours (p=4.1×10^−10^, q=39). They also showed clear discrimination between 1-hexanol (Hex Ret) and geraniol (Ger Ret) (p=4.6 × 10^−8^, q=29.87) showing that the covering and uncovering does not stress the bee and cause memory loss. **C)** *A dorsata* learned and retained memory after 3 hours with one antenna and two antenna. The acquisition and retention with two antenna were approximately same as would be predicted from the rates with one antenna if the olfactory pathway of each sided made decision independently (0.35*2-0.35*0.35=0.58 retention) and (0.38*2-0.38*0.38=0.62 acquisition) **D)** High acquisition (95%) and retention (0.75) rate in *Apis mellifera* even with one antenna, made two antenna case not very informative in this task. The retention to 1-Hexanol was seen to be stable once the trained ipsi antenna was unblocked at 3 hours. As mentioned in figure 3.B, discrimination between 1-hexanol and 1-geraniol was high even after coating and then removing the coat.

### The process of peeling away the paint does not shock the bee into forgetting

For n= 12 *Apis dorsata* we tested whether the coating and uncoating of the acrylic paint shock the bees into forgetting (Figure 5). To check this we first trained the bees with either one of the antenna blocked. 15 minutes before the retention test we coated the trained antenna with the paint, waited for it to dry, then uncovered the coat before testing for retention. The process of removing the coating did not cause the bees to forget the learnt information and memory retention was 99%.

### Bees that don’t show retention at 3hours on the untrained side do preserve it on the trained side

To further confirm that lack of memory on the untrained side seen in the trained *Apis mellifera* honey bees is not because of the loss of memory on the trained side, the same bees that were trained with one antenna and tested with the contra antenna were checked for trained antenna retention after removing the cover from the trained antenna (Figure 2 B, D). Retention upon carrying out this paradigm was 82% (n= 53) and significantly above both antenna closed group (p=4.1×10^−10^, q=39) (Figure 2 B). Moreover, the discrimination between the trained odor 1-Hexenol and the novel odor 1-Geraniol was significant (p=1.4×10^−8^, q=32.1).

### High learning rate in contra side when using a wall to separate the antennae

Bees were trained with a wall separating the antenna. Learning rate reached 90% For (N=10) progressively over the training. When tested, 50% learning was seen on the antenna on the other side of the wall in the 6^th^ trial itself (Fig. 3 A). In the same setup with a wall, even when the trained antenna was blocked with acrylic paint and the bees trained (Fig. 3 B) they learned gradually over the 5 trials. They attained a learning percentage of 65% (n=19) by the 5^th^ trial despite having the training antenna blocked. When the untrained antenna was tested in the 6^th^ trial the 65% learning was maintained. This fraction of bees retained the memory for 3 hours. Despite our best attempts it seemed impossible to robustly separate the two antennae with a wall.

**Figure 3:**
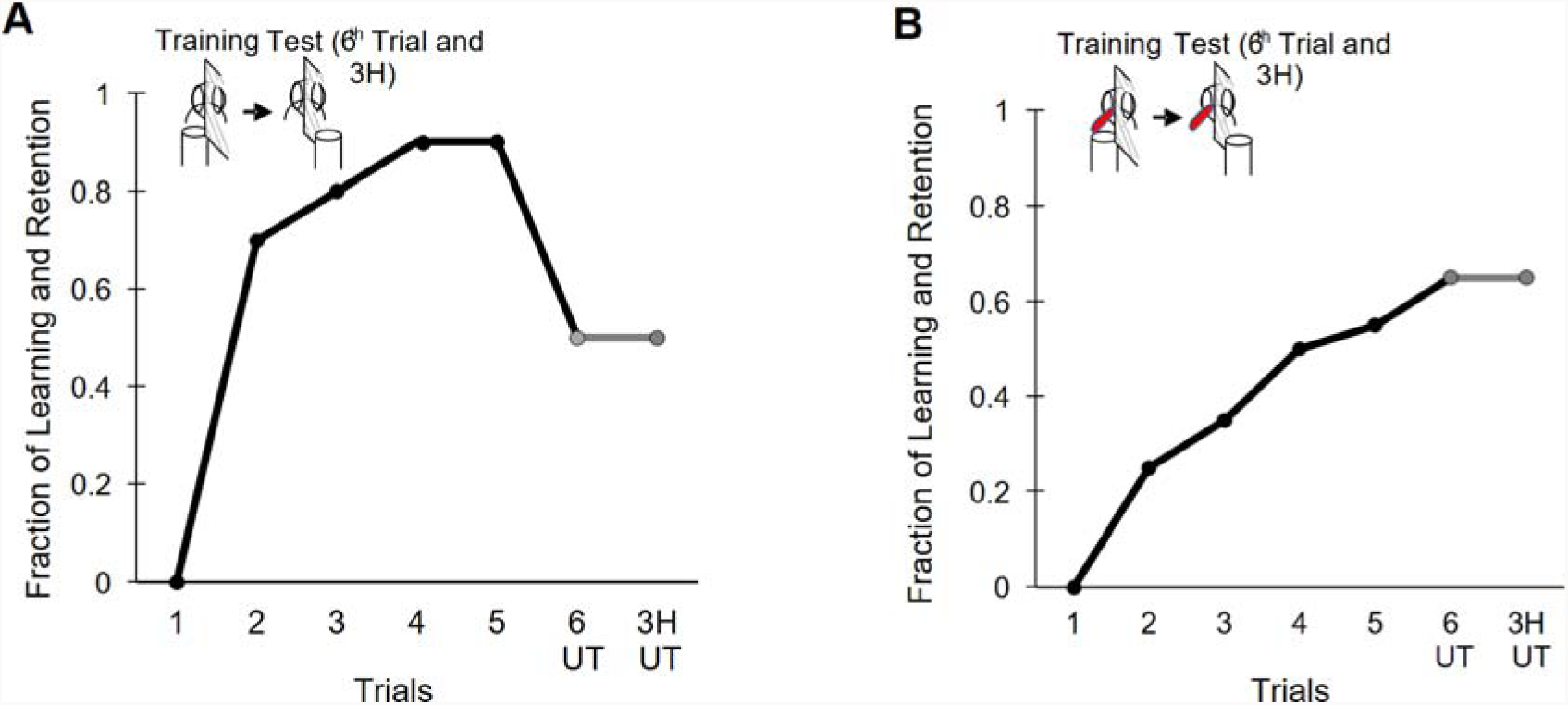
Training using a plastic wall to separate the two antennae shows learning in the contralateral side even as training is taking place. **A)** *Apis mellifera* exhibited PER to the trained odor on the untrained side at the 6^th^ trial itself when trained with plastic partition for isolation (denoted UT). This memory was retained on the untrained side at 3hrs post training (denoted 3 UT) (N=10). **B)** *Apis mellifera* exhibited learning on the trained side even when the trained antenna is insulated with acrylic and with plastic partition used for isolation. 65% retention was seen by the untrained antenna at the 6^th^ trial itself (denoted 6 UT), this memory was retained on the untrained side at 3hrs post training (denoted 3H UT) (N=19).

### The Acrylic paint block does not act as a contextual stimulus

Bees were trained with one antenna covered and split into two groups. One was tested without removing the coating and the other was tested with the coating removed. There is no observable difference in the percentage of retention between the bees with one antenna covered and the bees with the antennae uncovered at the time of testing (p=0.8, Cochran’s q= 0.05). No significant difference was seen in the discrimination either (p=0.1, Cochran’s q=2.6). The bees with their antenna uncovered discriminated marginally better than the bees with the one antenna covered (Fig. 4).

**Figure 4:**
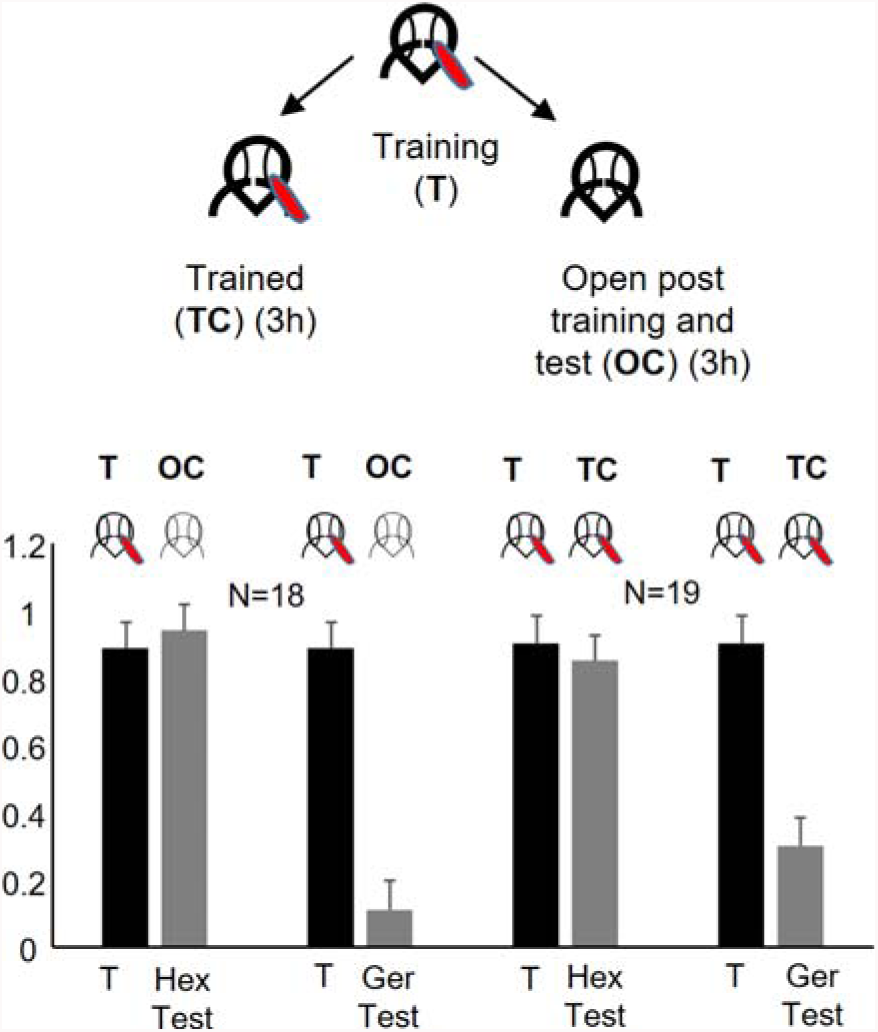
Bee does not learn the odor stimuli applied to one side alone as a stimulus different from odor presented to both sides: *Apis mellifera* bees were trained (T) with one antenna covered (N=37). At 3 hours the bees were divided randomly, in to two sets. A set of bees were checked for retention with the block opened (OC) (N=18) and the other set of bees were checked for retention in the trained condition (TC) (N=19). The bees with both antennae opened (OC) performed not differently with a 100% and 94% (TC) retention in each case. The bees with both antenna open and the bees with one antenna covered during retention test showed good discrimination (Ger Test OC and Ger Test TC).

**Figure 5:**
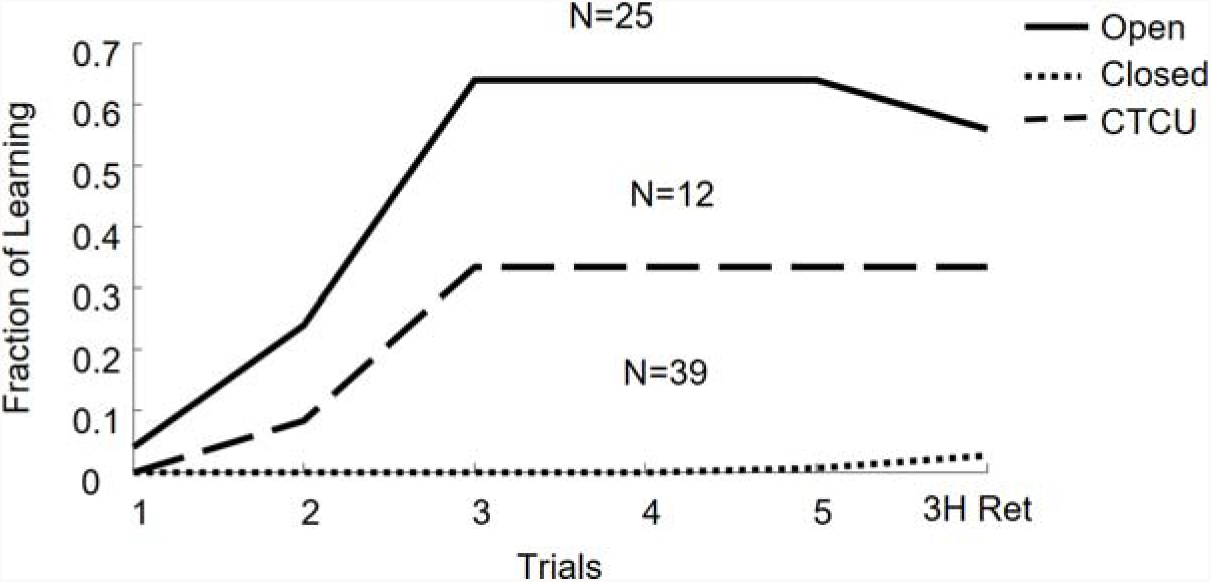
The process of coating and un-coating does not shock the bee into forgetting or harm the antenna. In *Apis dorsata* bees, the training antenna was coated with paint just prior to the 3 hour retention test (3h Ret, N=12). The removing of the coat does not shock the bees into forgetting the acquired memory as indicated by the 100% retention in the learnt bees.

## Discussion

Connections between the olfactory pathways of the two sides of the brain are prominent in lower and higher complex vertebrates in the Kingdom Animalia (Suarez et al., 2014). Bilateral connections have been shown in the visual system of insects (de Lussanet and Osse, 2012; Roper et al., 2017; Sanes and Zipursky, 2010). From the point of fundamental behaviour, the bilateral integration of vision would be advantageous especially for optimal orientation and direction alignment. For insects such as hymenopterans, olfaction is a dominant sense, imperative for the animal’s survival. However, the question remains, of how pivotal this bilateral integration and transfer of information is in other modalities like olfaction. Presence of a number of known bilateral neurons involved in PER associative conditioning with odor supports the possibility of transfer of association from one side of the brain to the other in the olfactory pathway. One multisensory mushroom body extrinsic neuron, the PE1, has been shown to display learning-related plasticity with respect to olfactory information in a time dependent manner (Mauelshagen, 1993; Menzel, 2012; Menzel and Benjamin, 2012). This neuron has it’s soma located ventro-medial to the α-lobe and arborizes adjacent to the contralateral α-lobe (Mauelshagen, 1993; Menzel and Muller, 1996; Okada et al., 2007). Given its anatomic positioning and learning dependant functional plasticity, the question about whether olfactory learned information with one antenna in honey bees can be retrieved from the contralateral side seemed a possibility. The ventral unpaired median neuron of the maxillary neuromere (VuMmx1) has it’s soma located at the subesophageal ganglion, its branches innervate, the basal lip of the MB, lateral horns, and antennal lobes, bilaterally (Hammer, 1993; Hammer, 1997; Hammer and Menzel, 1995). This bilateral neuron has also been shown to be octopaminergic positive and more crucially it displays plasticity upon olfactory learning (Hammer, 1993; Rein et al., 2013). This neuron can possibly act as the placeholder for reward bilaterally. However the bilateral transfer of olfactory memory is claimed to be odor specific and this would require either many neurons to use a population code or a very few neurons to use a complex temporal code. The evidence for using either of these by PE1 and VUM neurons is minimal. Thus it is not clear how the above mentioned neurons can be used to associate reward bilaterally in an odor specific way.

If we are to investigate the existence of lateral tracts in the olfactory pathway and mechanism of lateral we needed to first validate the existence of lateral transfer using a robust behavioural protocol. Given that this behavioural phenomena was said to have been observed in *Apis mellifera* (Sandoz and Menzel, 2001; Strube-Bloss et al., 2016) we attempted to observe the same behavioural output in a native Asian honey bee *A.dorsata*. However, over the course of our behavioural study, we did not observe lateral memory transfer and thus diminishing the possibility of finding such tracts in *A.dorsata* and these results remained consistent when we repeated the experiments using *Apis mellifera*.

### The learning and decision making in the two sides of in *Apis dorsata* are independent

We tested the hypothesis of whether the phenomenon of lateral transfer of olfactory information exists in *Apis dorsata.* Previous work on *Apis mellifera* suggested that when the two antennae were separated using a physical separator (plastic wall) and trained using one side, the learnt information could be retrieved using the contralateral untrained antenna (Sandoz and Menzel, 2001). Our result in *A.dorsata*, however, was quite contrary and not only did we see negligible transfer olfactory learnt information, but it was also observed that learning rate with one antenna in use, reduced significantly. We surmise that for this species of honey bee the learning is independent, implying the parallel working of both antennal lobes and olfactory pathways. In the same protocol, the *Apis mellifera* learning rate with one antenna reached up to 95% nearly saturating. In addition neuron tract tracing experiments from our lab using *Apis dorsata* showed no bilateral connections between the mushroom body calyx and the contralateral alpha lobe. Further, no connections were seen between the alpha lobe and contralateral antennal lobes (Mogily et al., 2018). These tract tracing experiments further strengthened the possibility of each lobe processing olfactory information independently. This is consistent with our result that the performance with both the antenna open case can be predicted using performance with one antenna, if independence of decision making on the two sides is assumed.

### Ecological significance of parallel pathways

The ecological importance of having parallel olfactory pathways in honey bees is still an enigma. Decision making for honey bees becomes crucial for each forager as the actions of the foragers eventually dictate the survival of the hive. Olfaction plays a massive role in the basic survivability of the honey bee as many of the decisions regarding food source would be made based on odor. Parallel systems can prove advantageous since the redundancy of the neural circuit would allow for optimum behaviour to manifest in the absence or loss of one of the two circuits, hence the bee will be able to perform its task despite the loss of function of one olfactory pathway. We do not have evidence to see that this happens often in nature. We hardly ever see bees with one antenna. The additive nature of this same circuit will prove beneficial as it would aid in improving the decision performance as seen in the experiments when we compare the performance of one antenna trained versus both trained case. Taking all this together, the results in both the species were consistent with no transfer of memory from the trained side to the untrained side and each side learning and retrieving independently. Our results remain consistent with the finding in 1972 by Mahsur et al., that with respect to olfaction the honey bee seems to use each lobe independently. It is not clear if there is a set of non-motor, decision neurons, that receive input from both the sides, or whether the two sides drive the motor neurons and thus the muscles independently and this requires further investigation. Our results question the possibility of finding robust odor coding bilateral tracts at higher level in honeybees.

## Supporting information

## Acknowledgement

We are grateful to the University Grants Commission India, UPE and DST-PURSE for funding our research. We are also grateful to CSIR for providing their support in funding the research via their fellowship and contingency for this work. We would like to thank the National Institute of Rural development and Panchayati Raj (NIRDPR Telengana) for providing us *Apis mellifera* honey bees. We would also like to thank Uttam Krishna Sharma and Sunil Kumar Sethy for their support in procuring and standardizing the collection of *Apis dorsata*.

## Paper References

1. Aboitiz. F., Montiel. J. (2003). One hundred million years of interhemispheric communication: the history of the corpus callosum. Braz. J. Med. Biol. Res. 36, 409–420.

2. Anfora, G., Rigosi, E., Frasnelli, E., Ruga, V., Trona, F., & Vallortigara, G. (2011). Lateralization in the invertebrate brain: left-right asymmetry of olfaction in bumble bee, Bombus terrestris. PLoS One, 6(4), e18903

3. Barron, A. B., & Klein, C. (2016). What insects can tell us about the origins of consciousness. Proc Natl Acad Sci U S A, 113(18), 4900–4908

4. Bitterman, M.E., Menzel, R., Fietz, A., and Schäfer, S. 1983. Classical conditioning of proboscis extension in honeybees (Apis mellifera). J. Comp. Psychol. 97: 107–119

5. Corballis, M. C. (2009). The evolution and genetics of cerebral asymmetry. Philos Trans R Soc Lond B Biol Sci, 364(1519), 867–879.

6. Corballis, M. C. (2017). The Evolution of Lateralized Brain Circuits. Front Psychol, 8, 1021.

7. de Lussanet, M. H. E., & Osse, J. W. M. (2012). An ancestral axial twist explains the contralateral forebrain and the optic chiasm in vertebrates. Animal Biology, 62(2), 193–216.

8. Frasnelli, E. (2013). Brain and behavioral lateralization in invertebrates. Front Psychol, 4, 939

9. Frasnelli, E., Haase, A., Rigosi, E., Anfora, G., Rogers, L. J., & Vallortigara, G. (2014). The Bee as a Model to Investigate Brain and Behavioural Asymmetries. Insects, 5(1), 120–138.

10. Gazzaniga, M. S. (2000). Cerebral specialization and interhemispheric communication: does the corpus callosum enable the human condition? Brain 123, 1293–1326.

11. Gazzaniga, M. S. (2014). The split-brain: rooting consciousness in biology. Proc Natl Acad Sci U S A, 111(51), 18093–18094.

12. Ghirlanda, S., Frasnelli, E., & Vallortigara, G. (2009). Intraspecific competition and coordination in the evolution of lateralization. Philos Trans R Soc Lond B Biol Sci, 364(1519), 861–866.

13. Gupta, N., & Stopfer, M. (2012). Functional analysis of a higher olfactory center, the lateral horn. J Neurosci, 32(24), 8138–8148.

14. Guo, Y., Wang, Z., Li, Y., Wei, G., Yuan, J., Sun, Y., … Chen, R. (2016). Lateralization of gene expression in the honeybee brain during olfactory learning. Sci Rep, 6, 34727.

15. Haase, A., Rigosi, E., Frasnelli, E., Trona, F., Tessarolo, F., Vinegoni, C., … Antolini, R. (2011). A multimodal approach for tracing lateralisation along the olfactory pathway in the honeybee through electrophysiological recordings, morpho-functional imaging, and behavioural studies. Eur Biophys J, 40(11), 1247–1258.

16. Hammer, M. (1993). An identified neuron mediates the unconditioned stimulus in associative olfactory learning in honeybees. Nature, 366, 59

17. Hammer, M. (1997). Neural basis of associative reward learning in honeybees. Trends Neurosci, (20) 245–252.

18. Hammer, M., Menzel, R (1995). Learning and memory in the honeybee. J Neurosci, 15(3), 1617–1630.

19. Hansson, B. S., & Stensmyr, M. C. (2011). Evolution of insect olfaction. Neuron, 72(5), 698–711.

20. J, Erber., T H, Masuhr., & R, Menzel. (1980). Localization of short-term memoy in the brain of the bee, Apis mellifera. Physiol Entomol, (5), 343–358.

21. Kaupp, U. B. (2010). Olfactory signalling in vertebrates and insects: differences and commonalities. Nat Rev Neurosci, 11(3), 188–200.

22. Kirschner, S., Kleineidam, C. J., Zube, C., Rybak, J., Grunewald, B., & Rossler, W. (2006). Dual olfactory pathway in the honeybee, Apis mellifera. J Comp Neurol, 499(6), 933–952.

23. Komischke, B., Sandoz, J. C., Malun, D., & Giurfa, M. (2005). Partial unilateral lesions of the mushroom bodies affect olfactory learning in honeybees Apis mellifera. Eur J Neurosci, 21(2), 477–485.

24. Letzkus, P., Ribi, W. A., Wood, J. T., Zhu, H., Zhang, S. W., & Srinivasan, M. V. (2006). Lateralization of olfaction in the honeybee Apis mellifera. Curr Biol, 16(14), 1471–1476.

25. Masuhr T., Menzel R. (1972) Learning Experiments on the Use of Side — Specific Information in the Olfactory and Visual System in the Honey Bee (Apis mellifica). In: Wehner R. (eds) Information Processing in the Visual Systems of Anthropods. Springer, Berlin, Heidelberg.

26. Mauelshagen J (1993) Neural correlates of olfactory learning in an identified neuron in the honey bee brain. J Neurophysiol 69:609–625.

27. Malun, D., Giurfa, M., Galizia, C. G., Plath, N., Brandt, R., Gerber, B., & Eisermann, B. (2002). Hydroxyurea-induced partial mushroom body ablation does not affect acquisition and retention of olfactory differential conditioning in honeybees. J Neurobiol, 53(3), 343–360.

28. Malun, D., Plath, N., Giurfa, M., Moseleit, A. D., & Muller, U. (2002). Hydroxyurea-induced partial mushroom body ablation in the honeybee Apis mellifera: volumetric analysis and quantitative protein determination. J Neurobiol, 50(1), 31–44.

29. Matsumoto, Y., Menzel, R., Sandoz, J.-C., & Giurfa, M. (2012). Revisiting olfactory classical conditioning of the proboscis extension response in honey bees: A step toward standardized procedures. Journal of Neuroscience Methods, 211(1), 159–167.

30. Menzel, R. (2012). The honeybee as a model for understanding the basis of cognition. Nat Rev Neurosci, 13(11), 758–768.

31. Menzel, R., & Benjamin, P R. (2012). In search of the engram in the Honeybee brain. Invertebrate Learning and Memory, Vol. 22, Chapter 29, pages (399–414).

32. Menzel, R., Muller, U. (1996). Learning and memory in honeybees: From behaviour to neural substrates. Annu rev neurosci (19), 379–404.

33. Mihrshahi, R. (2006). The corpus callosum as an evolutionary innovation. J Exp Zool B Mol Dev Evol, 306(1), 8–17.

34. Mogily, S., VijayKumar, M., Sethy, S. K., & Joseph, J. (2018). Characterization of the olfactory system in Apis dorsata, an Asian honey bee. bioRxiv. doi: 10.1101/420968.

35. Nawrot, M. P. (2012). Dynamics of sensory processing in the dual olfactory pathway of the honeybee. Apidologie, 43(3), 269–291.

36. Okada, R., Rybak, J., Manz, G., & Menzel, R. (2007). Learning-related plasticity in PE1 and other mushroom body-extrinsic neurons in the honeybee brain. J Neurosci, 27(43), 11736–11747.

37. Rein, J., Mustard, J. A., Strauch, M., Smith, B. H., & Galizia, C. G. (2013). Octopamine modulates activity of neural networks in the honey bee antennal lobe. Journal of Comparative Physiology. A, Neuroethology, Sensory, Neural, and Behavioral Physiology, 199(11), 947–962.

38. Roper, M., Fernando, C., & Chittka, L. (2017). Insect Bio-inspired Neural Network Provides New Evidence on How Simple Feature Detectors Can Enable Complex Visual Generalization and Stimulus Location Invariance in the Miniature Brain of Honeybees. PLoS Comput Biol, 13(2), e1005333.

39. Royet, J. P., & Plailly, J. (2004). Lateralization of olfactory processes. Chem Senses, 29(8), 731–745.

40. Rybak, J., Menzel, R. (1993). Anatomy of the MB in the honeybee brain the neuronal connections of the alpha lobe. J Comp Neurol, 334(3), 444–465.

41. Sandoz, J. C., Galizia, C. G., & Menzel, R. (2003). Side-specific olfactory conditioning leads to more specific odor representation between sides but not within sides in the honeybee antennal lobes. Neuroscience, 120(4), 1137–1148.

42. Sandoz, J. C., Hammer, M., & Menzel, R. (2002). Side-specificity of olfactory learning in the honeybee: US input side. Learn Mem, 9(5), 337–348.

43. Sandoz, J. C., & Menzel, R. (2001). Side-specificity of olfactory learning in the honeybee: generalization between odors and sides. Learn Mem, 8(5), 286–294.

44. Sanes, J. R., & Zipursky, S. L. (2010). Design principles of insect and vertebrate visual systems. Neuron, 66(1), 15–36.

45. Singh, S., & Joseph, J. (2018). Evolutionarily conserved anatomical and physiological properties of olfactory pathway till fourth order neurons in a species of grasshopper (Hieroglyphus banian). bioRxiv. doi: 10.1101/436626

46. Strausfeld, N. J. (2002). Organization of the honey bee mushroom body: representation of the calyx within the vertical and gamma lobes. J Comp Neurol, 450(1), 4–33.

47. Strube-Bloss, M., Nawrot, M. P., Menzel R. (2016). Neural correlates of side-specific odour memory in mushroom body output neurons. Proc. R. Soc. B 283: 20161270.

48. Su, C. Y., Menuz, K., & Carlson, J. R. (2009). Olfactory perception: receptors, cells, and circuits. Cell, 139(1), 45–59.

49. Suarez, R., Gobius, I., & Richards, L. J. (2014). Evolution and development of interhemispheric connections in the vertebrate forebrain. Front Hum Neurosci, 8, 497.

50. Vallortigara, G. (2006). The evolutionary psychology of left and right: costs and benefits of lateralization. Dev Psychobiol, 48(6), 418–427.

