## Supplementary material for "Evidence for absence of bilateral transfer of olfactory learned information in *Apis dorsata* and *Apis mellifera*"

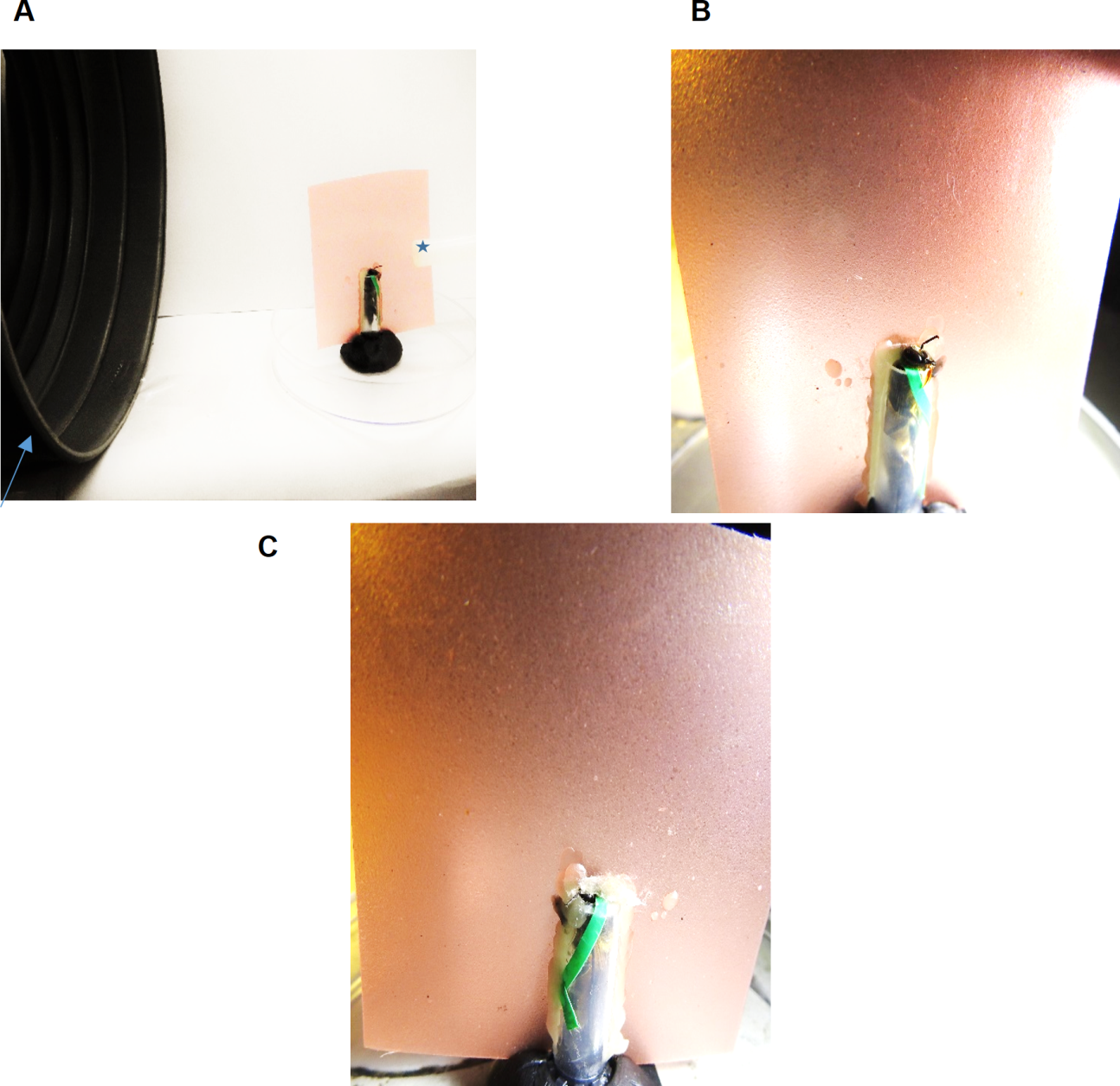

**Supplementary Figure 1: Setup for training with partition**, A) View of the setup with the 4mm x 5mm plastic wall odor exhaust (	) and odor valve (	)	B) Side View of training antenna C) Side view of untrained antenna.
